# A common Iba1 antibody labels vasopressin neurons in mice

**DOI:** 10.1101/2025.08.29.673171

**Authors:** Hannah D. Lichtenstein, Faith Kamau, Shaina McGrath, Javier E. Stern, Jessica L. Bolton

## Abstract

There are a wide variety of commercially available antibodies for labeling microglial cells based on different protein targets, as well as antibodies for the same protein target made in different species. While this array of targets and hosts allows for flexibility in immunohistochemical experiments, it is important to validate that different antibodies provide comparable and accurate immunodetection prior to experimental data collection. We found that a commercially available anti-Iba1 antibody, made in goat, produces irregular staining patterns in specific regions of the mouse brain, prompting a further investigation into the phenomenon. This Iba1-goat antibody displayed increased numbers of labeled cells when compared to expression of a CX3CR1-GFP reporter and IHC detection of P2RY12, two common microglial markers. Furthermore, immunodetection by other common anti-Iba1 antibodies made in rabbit and chicken did not display the excessive cell labeling when compared to the CX3CR1-GFP reporter. Upon further investigation, this Iba1-goat antibody was observed to highly colocalize with vasopressin neurons in the paraventricular nucleus of the hypothalamus (PVN) and the supraoptic nucleus of the hypothalamus (SON), the two main sites of vasopressin production in the brain. Other anti-Iba1 antibodies made in other species did not show this same colocalization with vasopressin. Finally, this effect was species-specific, as Wistar rats did not display erroneous cell labeling by the Iba1-goat antibody. In sum, the present study employs both qualitative and quantitative data to highlight the importance of validating antibody efficacy and specificity in a region- and species-specific manner.

**Significance Statement:** Microglia are the primary immune cells of the brain and are involved in many neurodevelopmental, as well as neurodegenerative, processes, thus making the study of microglia an important area of neuroscience research. There is a wide array of antibodies available to label microglia. Specific detection of microglia using immunohistochemistry is crucial for understanding differences in cell density, morphology, and interactions with other cells in various contexts. In the present study, a common anti-Iba1 antibody made in goat was found to display erroneous labeling of vasopressin neurons in specific regions of the mouse brain, inconsistent with other microglial markers, which emphasizes the importance of validating antibody efficacy and specificity in a region- and species-specific manner prior to beginning experimental data collection.

## Introduction

Microglia are the resident immune cells of the brain and are involved in many important neurodevelopmental and neurodegenerative processes (Bolton et al., 2022; Lenz et al., 2013; Miyamoto et al., 2016; Nelson et al., 2021; Paolicelli et al., 2011; Schafer et al., 2012). Thus, having the tools to visualize these cells in the brain are important for advancing neuroscience research, whether via transgenic expression of fluorescent reporters (for review, see Zhao et al., 2019) or immunohistochemistry (IHC; for review, see Korzhevskii & Kirik, 2016). Some of the most commonly used immunohistochemical targets for visualizing microglia are ionized calcium-binding adapter molecule 1 (Iba1), purinergic receptor P2Y12 (P2RY12), transmembrane protein 119 (TMEM119), and the fractalkine receptor (CX3CR1), all of which have varying levels of specificity for microglia vs. macrophages (for review, see Jurga et al., 2020).

Importantly, the signal intensity of these microglial markers can fluctuate depending on the age of an organism, due to the changes in gene expression within microglia throughout the lifespan (Hammond et al., 2019) (Gómez Morillas et al., 2021). Specificity is another factor that comes into play when choosing a microglial marker: while Iba1 may be the most popular target for visualizing microglia via IHC, this marker is also expressed by macrophages, whether brain-resident or infiltrating from the periphery (Ohsawa et al., 2004), which can sometimes localize quite close to microglia (Cronk et al., 2018).

Due to the nature of these tools and techniques, it is good scientific practice to use multiple markers for microglia when embarking on a research project that uses specific antibodies for the first time to validate the efficacy of any one antibody. Not only do different microglial markers have their unique limitations, issues with reproducibility in general can arise due to lot-to-lot variability, even when prepared from the same donor animal (Meliopoulos & Schultz-Cherry, 2018). Donor animals of different species, such as goat, rabbit, and chicken, allow for the use of multiple targets within one experiment, as antibodies developed within one species will typically not interfere with the binding of antibodies from other species. Thus, commonly used antibodies, such as Iba1, are typically manufactured in a variety of species, so that researchers are able to conduct experiments with different combinations of molecular targets in one tissue sample.

We recently observed that a common anti-Iba1 antibody, made in goat produced irregular staining patterns in select regions in the mouse brain, such as the paraventricular nucleus of the hypothalamus (PVN) and the supraoptic nucleus (SON): Microglia were labeled by Iba1, but so were cells that had a non-microglial morphology and colocalized with vasopressin. When alternative microglial markers were used, such as P2RY12 and CX3CR1-GFP, they did not colocalize with vasopressin. Furthermore, when different Iba1 antibody species were used, such as those made in chicken and rabbit, the unusual staining was not detected. The present study outlines the observed effects both qualitatively and quantitatively, highlighting the importance of validating antibody efficacy and specificity in a region- and species-specific manner prior to beginning experiments.

## Methods

### Animals

The mice used in this study were immature (postnatal day [P]8) or adult (P60-90) males and females from either a CX3CR1-GFP+/-reporter line (strain #005582, The Jackson Laboratory, RRID:IMSR_JAX:005582; where transgenic microglial reporters were included) or “wild-type” mice (i.e., Cre-/-) from a CX3CR1-BAC-Cre mouse line (MGI: 5311737; RRID: MMRRC: 036395-UCD), both from a C57BL/6 background (although the CX3CR1-GFP mice are the C57BL/6J substrain, whereas the CX3CR1-BAC-Cre mice are the C57BL/6N substrain). The rats used in this study were adult male Wistar rats (RccHan®:WIST, Envigo, RRID:RGD_13508588). Mice and rats were housed under 12 h light/dark cycle with free access to food and water. All experiments were performed in accordance with National Institutes of Health (NIH) guidelines and were approved by the [Author University] Animal Care and Use Committee.

### Experimental Design

The accuracy of the goat anti-Iba1 antibody was analyzed in CX3CR1-GFP+/- and wild-type mice. Animals were killed with Euthasol (Patterson Veterinary, VIRBAC Animal Health) and transcardially perfused with ice-cold 1x phosphate-buffer saline (PBS) followed by 4% paraformaldehyde. Perfused brains of CX3CR1-GFP+/-mice were then post-fixed in 4% paraformaldehyde in 0.1 M PBS for 4-6 h before cryopreserving in 15% sucrose solution overnight followed by 25% (for pups) or 30% (for adults) sucrose solution until brains sunk. Brains were frozen by dipping into 2-methylbutane for ∼30 s and were stored in a -80ºC freezer. Pups were weaned on P21 into standard, same-sex cages of 2–5 mice and killed with the same procedure as described before on P60.

### Immunohistochemistry (IHC)

Brains of CX3CR1-GFP+/- and wild-type mice were coronally sectioned into 25-µm-thick slices (1:4 series of the PVN and SON) using a Leica CM 1860 cryostat (RRID: SCR_025772) and stored in an anti-freeze solution in a -20ºC freezer. For IHC, floating brain sections were washed several times (3 x 5 min) with PBS-T (PBS containing 0.3% Triton X-100; Fisher Scientific Cat# AC215682500; pH=7.4) at room temperature (RT). Tissues were then allowed to permeabilize in 0.09% H_2_O_2_ in PBS-T for 20 min at RT. Sections were washed several times with PBS-T, then incubated in blocking solution containing 5% normal donkey serum (NDS; Jackson ImmunoResearch Cat# 017-000-121, RRID: 2337258) for 1 h at RT to prevent non-specific binding. Sections were then incubated in primary antibody solution overnight at 4ºC. The next morning, sections were washed in PBS-T several times and incubated in secondary antibody solution (see Table 1) for 3 h at RT. After repeated PBS-T washes, free-floating sections were counter-stained with DAPI for 1 min before mounting on gelatin-coated slides and coverslipping with Fluormount-G Mounting Medium (Thermo Fisher Scientific Cat# 00-4958-02, RRID: SCR_015961). Immunostaining P8 brain sections with anti-Iba1-chicken and anti-Iba1-rabbit required antigen-retrieval with Tris-EDTA buffer at 90ºC for 3 min prior to the blocking step before continuing with the protocol described above.

**Table 1:** Summary of antibodies used for immunohistochemical experiments. From left to right: Primary antibody, associated concentration and buffer, secondary antibody, associated concentration and buffer.

| Primary antibody | Concentration & Buffer | Secondary antibody | Concentration & Buffer |
| --- | --- | --- | --- |
| Goat anti-Iba1 (Iba1-goat), FujiFilm WAKO Pure Chemical Corporation Cat# 011-27991, RRID: AB_2935833<br>Lots: SKK1868, LEG4278, WTQ1615 | 1:1000<br>2% NDS,<br>0.3% Triton<br>X-100 | Donkey anti-goat IgG-568, Thermo Fisher Scientific Cat# A11057, RRID: AB_2534104, Lot #23044269<br>Donkey anti-Goat IgG-647, Jackson ImmunoResearch Lab Cat# 705-607-003, RRID: AB_2340439 | 1:500<br>2% NDS,<br>0.3% Triton<br>X-100 |
| Chicken anti-Iba1 (Iba1-chicken), Synaptic Systems Cat# 234-009, RRID: AB_2891282<br>Lots: 1-11, 1-16 | 1:2000<br>2% NDS,<br>0.3% Triton<br>X-100 | Alexa Fluor-568 AffiniPure Donkey anti-Chicken IgG, Thermo Fisher Scientific Cat# A78950, RRID: AB_291072, Lot #2622381<br>Alexa Fluor-647 Donkey anti-Chicken IgG, Jackson ImmunoResearch Labs Cat# 703-605-155, RRID: AB_2340379, Lots #160355, 2540901 | 1:1000<br>2% NDS,<br>0.3% Triton<br>X-100 |
| Rabbit anti-Iba1 (Iba1-rabbit), FUJIFILM Wako Pure Chemical Corporation Cat# 019-19741, RRID: AB_839504<br>Lot: SEP6150 | 1:2000<br>2% NDS,<br>0.3% Triton<br>X-100 | Donkey anti-Rabbit IgG, Alexa Fluor 568, Thermo Fisher Scientific Cat# A10042, RRID: AB_2534017, Lots #2306809, 2540901 | 1:1000<br>2% NDS,<br>0.3% Triton<br>X-100 |
| Rabbit anti-P2RY12, AnaSpec; EGT Group Cat#AS-55043A, RRID: AB_2298886<br>Lots: WF0501, WTQ1615 | 1:2000<br>2% NDS,<br>0.3% Triton<br>X-100 | Alexa Fluor 647-AffiniPure Donkey anti-Rabbit IgG, Jackson ImmunoResearch Labs Cat# 711-605-152, RRID: AB_2492288, Lots# 167518, 173915 | 1:1000<br>2% NDS,<br>0.3% Triton<br>X-100 |
| Rabbit anti-Vasopressin-Neurophysin (VP-NP), Dr. Alan Robinson, University of Pittsburgh Cat# rn2, RRID: AB_2751000 | 1:20,000<br>2% NDS,<br>0.3% Triton<br>X-100 | Donkey anti-Rabbit IgG, Alexa Fluor 568, Thermo Fisher Scientific Cat# A10042, RRID: AB_2534017, Lots #2306809, 2540901<br>Donkey anti-Rabbit IgG, Alexa Fluor 488, Thermo Fisher Scientific Cat# A-21206, RRID: AB_2535792, Lots #2289872, 2873188 | 1:500<br>2% NDS,<br>0.3% Triton<br>X-100 |
| Mouse anti-Oxytocin-Neurophysin (OT-NP), Ben-Barak et al., 1985, Cat# PS36, RRID: AB_2315027 | 1:20,000<br>2% NDS,<br>0.3% Triton<br>X-100 | Donkey anti-Mouse IgG, Alexa Fluor 568, Thermo Fisher Scientific Cat# A10037, RRID: AB_1180865, Lot #2555709 | 1:500<br>2% NDS,<br>0.3% Triton<br>X-100 |

Immunostaining for vasopressin-neurophysin (VP-NP; Verbalis & Robinson, 1983) and oxytocin-neurophysin (OT-NP; Ben-Barak et al., 1985) expression differed slightly from the protocol described above. After incubation in blocking solution of 5% NDS for 1 h at RT to prevent non-specific binding, sections were incubated with the primary antibody solution for anti-VP-NP-rabbit and/or anti-OT-NP-mouse overnight at 4ºC. The next day, sections were washed in PBS-T several times and incubated in secondary antibody solution (see Table 1) for 3 h at RT. After repeated PBS-T washes, tissues were incubated in blocking solution containing 5% NDS for 1 h at RT. Sections were then incubated with the primary antibody solution for anti-Iba1-goat overnight. The next day, sections were washed several times with PBS-T, then incubated with the secondary antibody solution (see Table 1) for 3 h at RT. The protocol then continues as described above. Methodological details, Cat#, and RRID for each antibody are specified in Table 1.

### Analysis

Images for the comparison of goat, chicken, and rabbit anti-Iba1 antibodies were taken with an Olympus BX41 Epifluorescent Clinical Microscope (RRID: SCR_027355). All other images were collected using a Zeiss LSM-780 Confocal Laser Scanning Microscope (RRID: SCR_020922) with a 20x objective. 13 Z-stack images were acquired at 1-µm intervals. The image frame was digitized at 16-bit using a 1024 x 1024-pixel frame size. Images for the Wistar rats used in Figure 5 were collected using a Zeiss LSM-980 with Airyscan 2 Microscope (RRID:SCR_025048) with a 40x objective. 70 Z-stack images were acquired at 0.35-µm intervals. The image frame was digitized at 16-bit using a 1024 x 1024-pixel frame size. Using Fiji (RRID: SCR_002285), an ROI was manually drawn around the PVN or SON, then the number and co-localization of Iba1+ microglia, P2RY12+ microglia, VP+, OT+ and GFP-reporter+ microglia were counted manually. Statistical differences were assessed using GraphPad Prism software (RRID: SCR_002798). Each group consisted of n=3 animals (comprised of both sexes). Two-way ANOVA tests were used to analyze the accuracy of the goat anti-Iba1 antibody using colocalization with the different microglial markers as the independent variables. Significant interactions were followed by Šídák post hoc tests. The significance level was set to 0.05 for all tests, and data are represented as mean ± SEM. All experiments were performed blindly without prior knowledge of the experimental group.

## Results

### Iba1-goat antibody displays increased cell-labeling compared to P2RY12 and CX3CR1-GFP expression, in both pups and adult mice, in the PVN and SON

Using CX3CR1-GFP+/- mice that have transgenic fluorescent reporters in microglia, we examined the difference in density of cell-labeling compared to Iba1-goat and P2RY12, two microglial markers commonly used in IHC, in both pups (Fig.1A,B) and adult mice (Fig.1D,E). Across pups and adults in both the PVN and SON, expression of CX3CR1-GFP and P2RY12 showed similar densities of labeled cells, whereas Iba1-goat resulted in a significantly increased density of labeled cells. In pups, there was a trending increase in the density of Iba1-goat+ cells compared to CX3CR1-GFP+ cells in the PVN (Fig.1A,C), and a significant increase in the density of Iba1-goat-labeled cells compared to both CX3CR1-GFP+ and P2RY12+ cells in the SON (Fig.1B,C). These results suggest that the effect may be stronger in the SON than PVN at this age. In adults, both the PVN and SON had a significant increase in the density of Iba1-goat+ cells compared to both CX3CR1-GFP+ and P2RY12+ cells (Fig.1D,F), suggesting a more prominent effect in adulthood. Interestingly, this effect was only evident in the PVN and SON, as other brain regions like the hippocampus and cortex, did not display excessive cell labeling (Fig.1-1).

### Iba1-goat displays excessive labeling of non-microglial cells compared to Iba1-rabbit and Iba1-chicken, in both pups and adult mice, in the PVN and SON

Using the same CX3CR1-GFP+/- mice, we next examined how three different versions of the Iba1 antibody, made in various host species, labeled microglia when compared to CX3CR1-GFP (Fig.2A,D). There was a significant increase in the density of Iba1-goat+ cells in the PVN of pups when compared with other Iba1 species (Fig.2B) and compared to colocalization with CX3CR1-GFP (Fig. 2B). In the SON of pups, there was a significant increase in the density of Iba1-goat+ cells when compared with Iba1-rabbit (Fig.2C) or Iba1-chicken (Fig.2C). The density of Iba1-goat+ cells was significantly increased when compared to colocalization with CX3CR1-GFP (Fig.2C), whereas Iba1-chicken+ cells was significantly decreased when compared to colocalization with CX3CR1-GFP (Fig.2C). In the PVN of adults, there was a significant increase in the density Iba1-goat+ cells when compared with other species of anti-Iba1 (Fig.2E) and compared to colocalization with CX3CR1-GFP (Fig.2E). Iba1-chicken displayed increased colocalization with CX3CR1-GFP+ cells compared with only Iba1+ cells (Fig.2E) and compared with Iba1-goat colocalization with CX3CR1-GFP (Fig.2E). In the SON of adults, there was a significant increase in the density of Iba1-goat+ cells when compared with other Iba1 species (Fig.2F) and compared to colocalization with CX3CR1-GFP (Fig.2F). These findings show that Iba1-goat labels microglial cells and many other cells that are not CX3CR1-GFP+, in both the PVN and SON of pups and adults, suggesting that the Iba1-goat antibody has off-target labeling in certain regions of the mouse brain.

### Iba1-goat displays high colocalization with vasopressin, and to some degree oxytocin, in the PVN and SON of adult mice

In order to better understand the identity of the non-microglial cells that are labeled by Iba1-goat, we examined the morphology of the cells and postulated that the Iba1-goat+, non-microglial cells appeared to look quite similar to that of principal neurons. Since the localization of this excessive Iba1-goat signal is so strong in the PVN and SON, and these are the main brain regions that produce vasopressin (VP) and oxytocin (OT), we investigated whether or not Iba1-goat colocalized with either of these signals in wild-type C57BL/6 mice (Fig.3A-B). When comparing the percentage of Iba1+ cells labeled by Iba1-goat alone, Iba1-goat colocalized with VP, and Iba1-goat colocalized with OT (Fig.3C), we found that ∼40% and 55% of the Iba1-labeled cells in the PVN and the SON, respectively, are co-labeled with VP. In the PVN, ∼20% of the Iba1+ cells are co-labeled with OT, whereas in the SON, less than 10% of the Iba1+ cells are co-labeled with OT. In the PVN, the remaining Iba1+ cells were labeled with Iba1-goat only, making up ∼25% of the Iba1-labeled cells, compared to the SON, wherein ∼50% of the Iba1+ cells were labeled with Iba1-goat only, likely indicating the number of true microglia/macrophages. These results suggest that although the Iba1-goat antibody does label microglia, it also shows off-target effects, labeling many vasopressin- and oxytocin-expressing neurons in the PVN and SON.

### When compared to Iba1-goat, Iba1-chicken displays little to no colocalization with vasopressin, and high levels of colocalization with CX3CR1-GFP

After observing considerable colocalization of Iba1-goat with VP, we wanted to confirm whether or not this pattern was observed with Iba1-chicken (Fig.4A-B). Using CX3CR1-GFP +/-mice, we confirmed that Iba1-goat displayed elevated co-labeling with VP in the PVN (∼66% of cells) and SON (∼37% of cells), with only ∼10-12% of Iba1-goat+ cells co-labeling with CX3CR1-GFP expression in both the PVN and SON, and with the remainder of the Iba1+ cells in the PVN (∼21%) and SON (∼50%) labeling only with Iba1-goat (Fig.4C). In contrast, using the same subjects, Iba1-chicken displayed a different pattern of colocalization with these markers: <5% of the cells labeled by Iba1-chicken colocalized with VP in both the PVN and SON, and ∼88% and 93% of the Iba1-chicken-labeled cells colocalized with CX3CR1-GFP, in the PVN and SON, respectively (Fig.4D). The remaining cells that were labeled by only Iba1-chicken made up ∼10% and 5% of the total cells labeled by Iba1-chicken in the PVN and SON, respectively. These remaining cells labeled with only Iba1 and not co-labeled with CX3CR1-GFP or VP may be labeling barrier-associated macrophages, and in the case of Iba1-goat, also OT cells.

### The irregular staining patterns of Iba1-goat are species-specific and are not observed in Wistar rats

In order to understand if this irregular staining pattern of Iba1-goat is specific to mice, we used samples from adult Wistar rats to perform IHC for anti-Iba1-goat and VP. Interestingly, in both the PVN and SON of rats, VP staining and anti-Iba1-goat staining were observed to be normal (Fig.5). These qualitative results suggest that the anti-Iba1-goat antibody (FujiFilm WAKO Pure Chemical Corporation Cat# 011-27991, RRID: AB_2935833) displays irregular staining in a species-specific manner.

## Discussion

The goal of this manuscript was to evaluate the efficacy of microglial cell-labeling by 3 commonly used anti-Iba1 antibodies made in 3 separate host species when compared with other conventional microglial markers, i.e., anti-P2RY12 antibody and CX3CR1-GFP reporter. A secondary goal of this manuscript was to raise awareness of the importance of antibody validation in scientific research. The validation of antibodies is crucial prior to experimental procedures in order to avoid erroneous results and misinterpretation of scientific phenomenon. Furthermore, it is important to validate antibody efficacy when using different lots of antibodies in order to ensure that results are consistent across batches. Although anti-Iba1 is one of the most commonly used antibodies for labeling microglial cells, we find in this study that the use of a specific version of this antibody made in goat, across multiple different lots (RRID: AB_2935833; lots SKK1868, LEG4278, WTQ1615), displays erroneous cell labeling of non-microglial cells in two brain regions, the PVN and the SON. These regions are also the primary source of vasopressin and oxytocin production, and further investigation into this phenomenon revealed that the erroneous cell-labeling colocalized most highly with vasopressin neurons in these regions.

While multiple recent studies have used anti-Iba1-goat in C57BL/6 mice and did not report this erroneous cell labeling (Cao et al., 2024; Hampton et al., 2020; Iguchi et al., 2023; Lee et al., 2025; Soares et al., 2025; Wang et al., 2025), these papers mostly focused on hippocampal and cortical brain regions (which exhibited normal staining in our hands as well, see Fig. 1-1), and did not investigate microglia within the PVN and SON regions of the hypothalamus using this antibody. Furthermore, it is possible that any lab that has previously attempted to utilize this anti-Iba1-goat antibody and found strange staining in the PVN and SON pivoted to use anti-Iba1 made in another host species or another microglial marker entirely, which is perhaps why this phenomenon has not been reported previously. While this antibody seems to have efficacy issues in mouse tissue in these regions, these issues do not arise in Wistar rats (Fig. 5), suggesting that the cross-reactivity of this antibody with vasopressin-expressing principal neurons in the PVN and SON is specific to mice. While the goal of this manuscript was not to investigate the reason why this anti-Iba1-goat antibody seems to have cross-reactivity with vasopressin-expressing cells in the PVN and SON of C57BL/6 mice, this mechanism would be interesting to investigate in future work, as well as whether this phenomenon occurs in other mouse strains.

In order to understand if the observations of erroneous anti-Iba1-goat staining was a developmental phenomenon or was consistent across the lifespan, both pups and adults of both sexes were used in the first two experiments when comparing this anti-Iba1-goat antibody to alternative microglial markers (Fig. 1) or with other anti-Iba1 antibodies made in other host species (Fig. 2). In both cases, and in both the PVN and the SON, there were clear increases in the densities of cell-labeling by anti-Iba1-goat compared to other microglial markers. These qualitative disparities were substantiated by our quantitative data, supporting the idea that this anti-Iba1-goat antibody labels more than just microglial cells in the PVN and SON across the lifespan in C57BL/6 mice.

**Figure 1.**
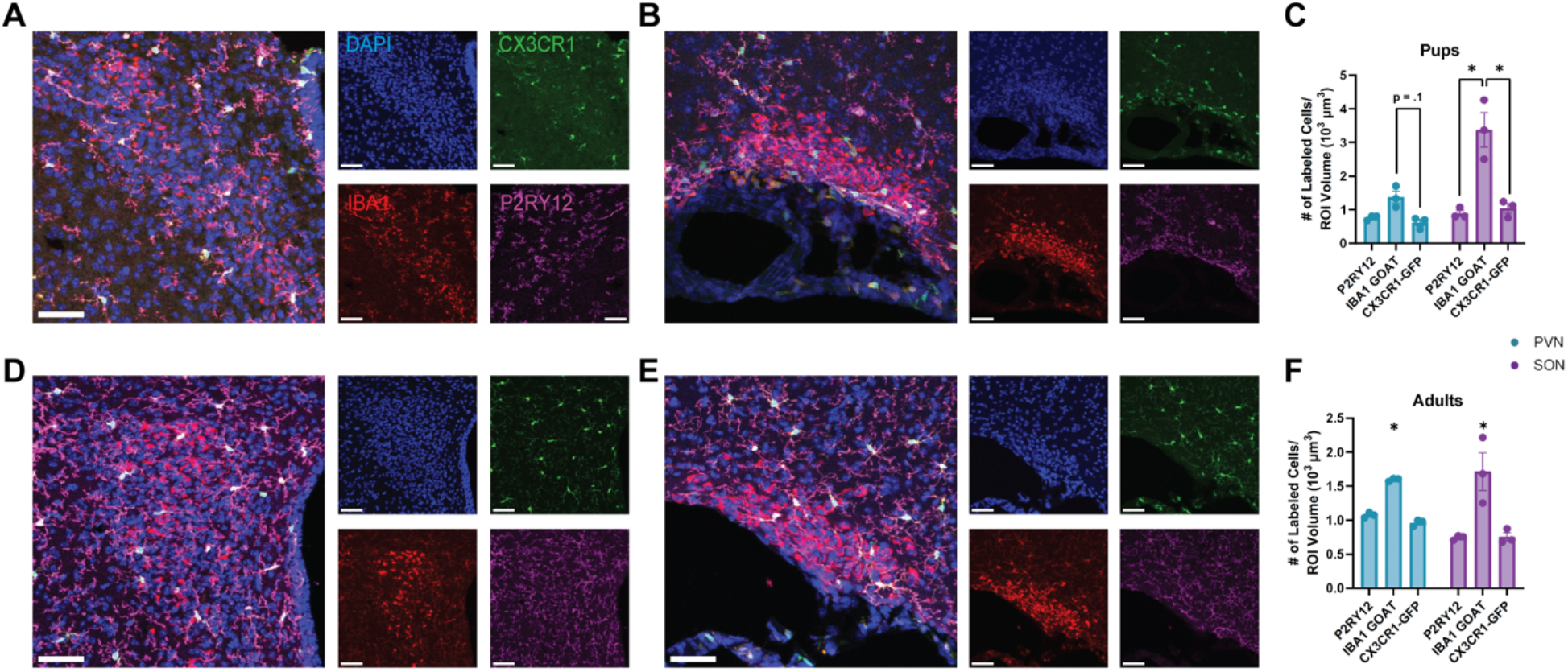
Iba1-goat displays increased cell labeling compared to P2RY12 and CX3CR1-GFP expression, in both pups and adult mice, in the PVN and SON. A-B) Representative images of the PVN (A) and SON (B) of pups. C) In the PVN, Iba1-goat antibody displayed a slight increase in the number of cells labeled compared to CX3CR1-GFP, and in the SON, Iba1-goat displayed a significant increase in the number of cells labeled within the region of interest compared to both P2RY12 and CX3CR1-GFP *(significant Brain Region x Microglial Marker interaction; F(2,4) = 14*.*32, p = 0*.*02; post-hoc tests, p*_*PVN Iba1-goat vs. CX3CR1*_ *= 0*.*1, p*_*PVN Iba1-goat vs. P2RY12*_ *= 0*.*2, p*_*SON Iba1-goat vs. P2RY12*_ *= 0*.*002, p*_*SON Iba1-goat vs. CX3CR1-GFP*_ *= 0*.*002)*. D-E) Representative images of the PVN (D) and SON (E) of adults. F) In the PVN and SON, Iba1-goat antibody displayed an increase in the number of cells labeled compared to both P2RY12 and CX3CR1-GFP *(significant main effect of Microglial Marker; F(2,8) = 23*.*45, p = 0*.*0005)*. Data are Mean ± SEM; *p<0.05. Scale bar = 20 μm, blue = DAPI, green = CX3CR1-GFP, red = Iba1-goat, magenta = P2RY12.

**Figure 2.**
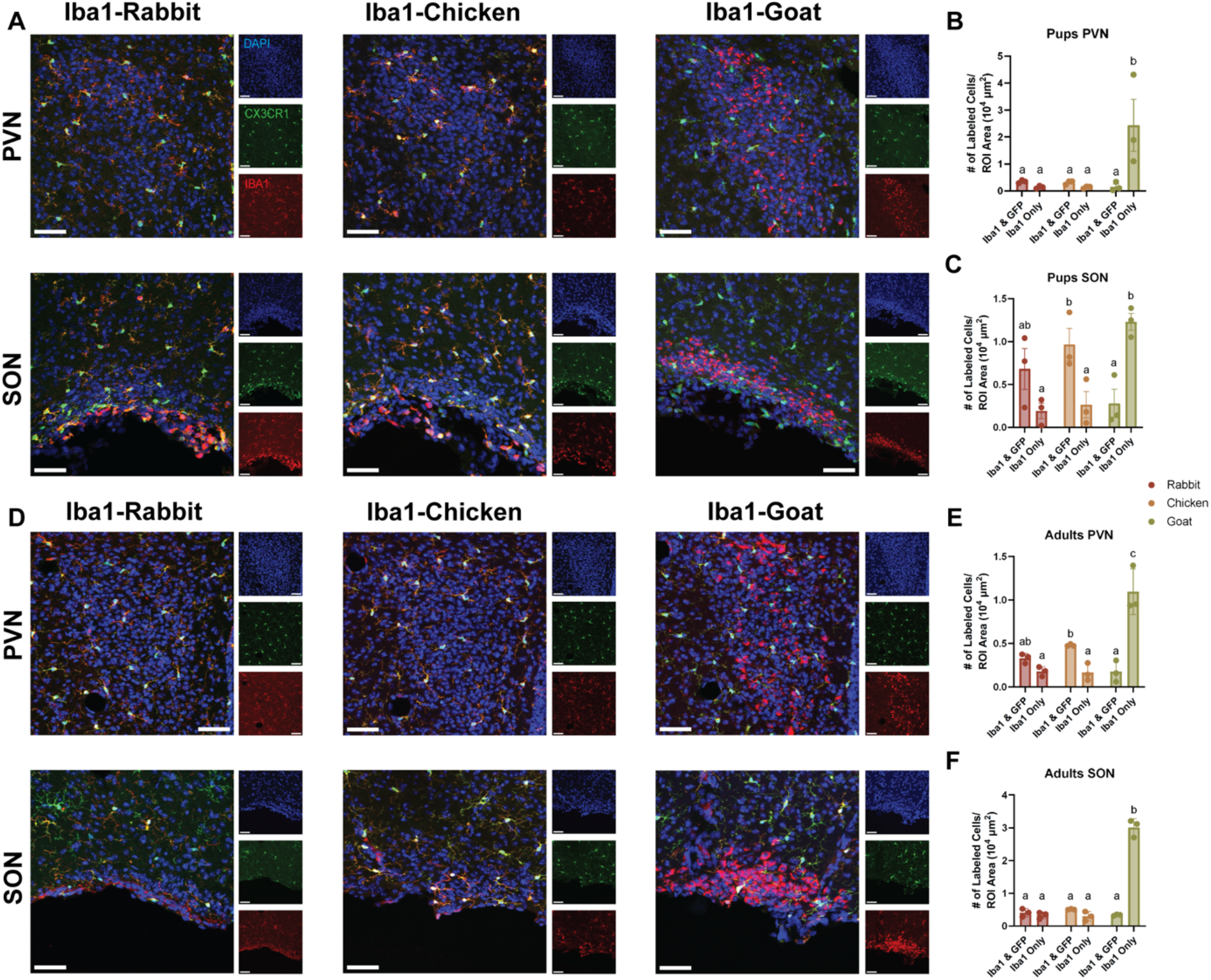
Iba1-goat displays excessive labeling of non-microglial cells compared to Iba1-rabbit and Iba1-chicken, in both pups and adult mice, in the PVN and SON. (A, D) Representative images, from left to right, top to bottom, of Iba1-rabbit, Iba1-chicken, and Iba1-goat antibodies immunostaining in the PVN and SON in pups (A) and adults (D). B) In the PVN of pups, Iba1-goat antibody displayed an increase in the number of cells labeled compared to Iba1-goat antibody colocalized with CX3CR1-GFP within the same tissue, as well as when compared to the number of cells labeled with Iba1-chicken and Iba1-rabbit *(significant Cell Density x Iba1 Species interaction; F(2,6) = 7*.*996, p = 0*.*02; post-hoc test, p*_*Iba1 only, rabbit vs. goat*_ *= 0*.*005*^*b*^, *p*_*Iba1 only, chicken vs. goat*_ *= 0*.*005*^*b*^, *p*_*Iba1-goat, Iba1 only vs. Iba1&GFP*_ *= 0*.*004*^*b*^). C) In the SON of pups, Iba1-goat antibody displayed an increase in the number of cells labeled compared to Iba1-goat & CX3CR1-GFP colocalized within the same tissue, as well as when compared to the number of cells labeled with Iba1-chicken and Iba1-rabbit *(significant Cell Density x Iba1 Species interaction; F(2,6) = 17*.*40, p = 0*.*003; post-hoc test, p*_*Iba1 only, rabbit vs. goat*_ *= 0*.*002*^*b*^, *p*_*Iba1 only, chicken vs. goat*_ *= 0*.*003*^*b*^, *p*_*Iba1-goat, Iba1 only vs. Iba1&GFP*_ *= 0*.*005*^*b*^, *p*_*Iba1 & GFP, chicken vs. goat*_ *= 0*.*03*^*a*^, *p*_*chicken, Iba1 only vs. Iba1&GFP*_ *= 0*.*02*^*a*^). E) In the PVN of adults, Iba1-goat antibody displayed an increase in the number of cells labeled compared to Iba1-goat & CX3CR1-GFP colocalized within the same tissue, as well as when compared to the number of cells labeled with Iba1-chicken and Iba1-rabbit *(significant Cell Density x Iba1 Species interaction; F(2,6) = 93*.*24, p < 0*.*0001; post-hoc test, p*_*Iba1 only, rabbit vs. goat*_ *< 0*.*0001*^*c*^, *p*_*Iba1 only, chicken v goat*_ *< 0*.*0001*^*c*^, *p*_*Iba1-goat, Iba1 only vs. Iba1&GFP*_ *<0*.*0001*^*c*^, *p*_*chicken, Iba1 only vs. Iba1&GFP*_ *= 0*.*004*^*b*^, *p*_*Iba1 & GFP, chicken vs. goat*_ *= 0*.*04*^*b*^). F) In the SON of adults, Iba1-goat antibody displayed an increase in the number of cells labeled within the region of interest compared to Iba1-goat & CX3CR1-GFP colocalized within the same tissue, as well as when compared to the number of cells labeled with Iba1-chicken and Iba1-rabbit *(significant Cell Density x Iba1 Species interaction; F(2,6) = 176*.*44, p < 0*.*0001; post-hoc test, p*_*Iba1 only, rabbit vs. goat*_ *< 0*.*0001*^*b*^, *p*_*Iba1 only, chicken vs. goat*_ *< 0*.*0001*^*b*^, *p*_*Iba1-goat, Iba1 only vs. Iba1&GFP*_ *< 0*.*0001*^*b*^). Data are Mean ± SEM; Scale bar = 20 μm, blue = DAPI, green = CX3CR1, red = Iba1 from either rabbit (left), chicken (center), or goat (right).

Interestingly, we observed some slight differences in cell-labeling densities amongst the other anti-Iba1 species: in the SON (but not the PVN) of pups, Iba1-chicken displays a significant increase in the levels of colocalization with CX3CR1-GFP compared to the other anti-Iba1 species (Fig. 2C), and the same phenomenon was observed in the PVN of adult mice (but not in the SON). This suggests that, in some cases, anti-Iba1-chicken may provide the most similar cell-labeling results to CX3CR1-GFP, a transgenic reporter for microglia.

Importantly, when considering the specificity of Iba1 compared to CX3CR1 in brain tissue, Iba1 labels monocytes/macrophages generally, whereas CX3CR1 is more highly expressed by microglia compared to perivascular macrophages (Mills et al., 2025; Mondo et al., 2020). When comparing the colocalization of Iba1-goat or Iba1-chicken with CX3CR1-GFP or VP, we found that there were a portion of cells in both the PVN and SON that have only Iba1 labeling (Fig. 4). In the context of Iba1-goat, this may be predominately due to partial colocalization with oxytocin (Fig. 3C), but this staining may also represent peripheral macrophages, especially as the regions analyzed in this study are bordering the ventricles. It has been previously shown that Iba1 primarily labels microglia, but also labels other myeloid cells such as macrophages (Ito et al., 1998; Kenkhuis et al., 2022; Unger et al., 2018), which could explain the 5-10% of Iba1-chicken+ cells that did not colocalize with CX3CR1-GFP in our data (Fig. 4D).

**Figure 3.**
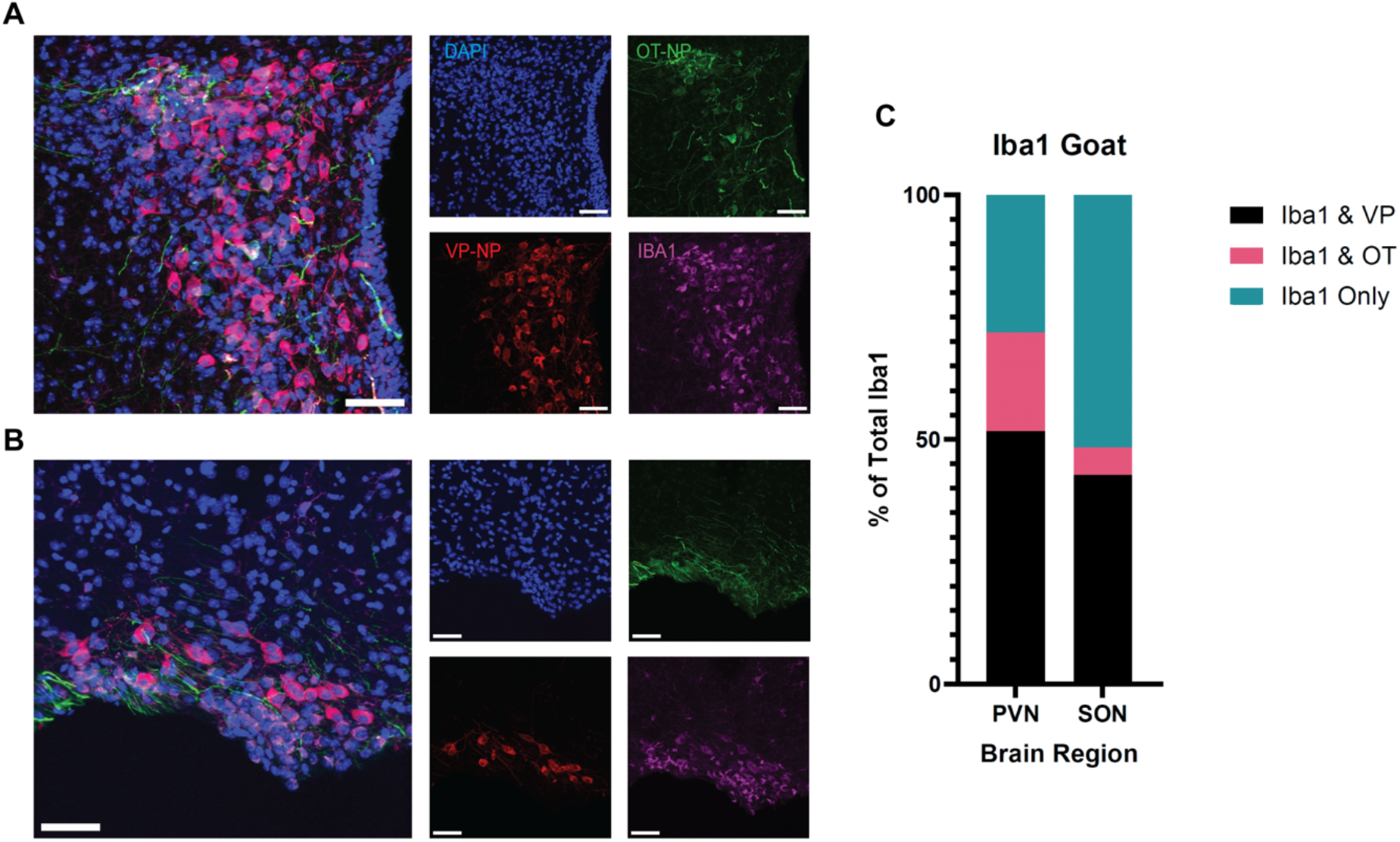
Iba1-goat displays high colocalization with vasopressin, and to some degree oxytocin, in the PVN and SON of adult mice. A-B) Representative images of the PVN (A) and SON (B). C) Comparison of the percentage of cells labeled by Iba1-goat antibody alone, Iba1-goat colocalized with VP, and Iba1-goat colocalized with OT within the PVN and the SON. 51.7%±13.3 of cells in the PVN and 42.7%±2.9 of cells in the SON were co-labeled with Iba1-goat and VP. In the PVN, 20.2%±12.6 of the cells were co-labeled with Iba1-goat and OT, whereas in the SON, 5.6%±0.7 of the cells were co-labeled with Iba1-Goat and OT. In the PVN, the remaining cells were labeled with Iba1-goat antibody only, making up 28.1%±0.8 of the labeled cells, as opposed to the SON, wherein 51.7%±2.2 of the cells were labeled with Iba1-goat only. Data are Mean ± SEM. Scale bar = 20 μm, blue = DAPI, green = VP, red = OT, magenta = Iba1-goat.

**Figure 4.**
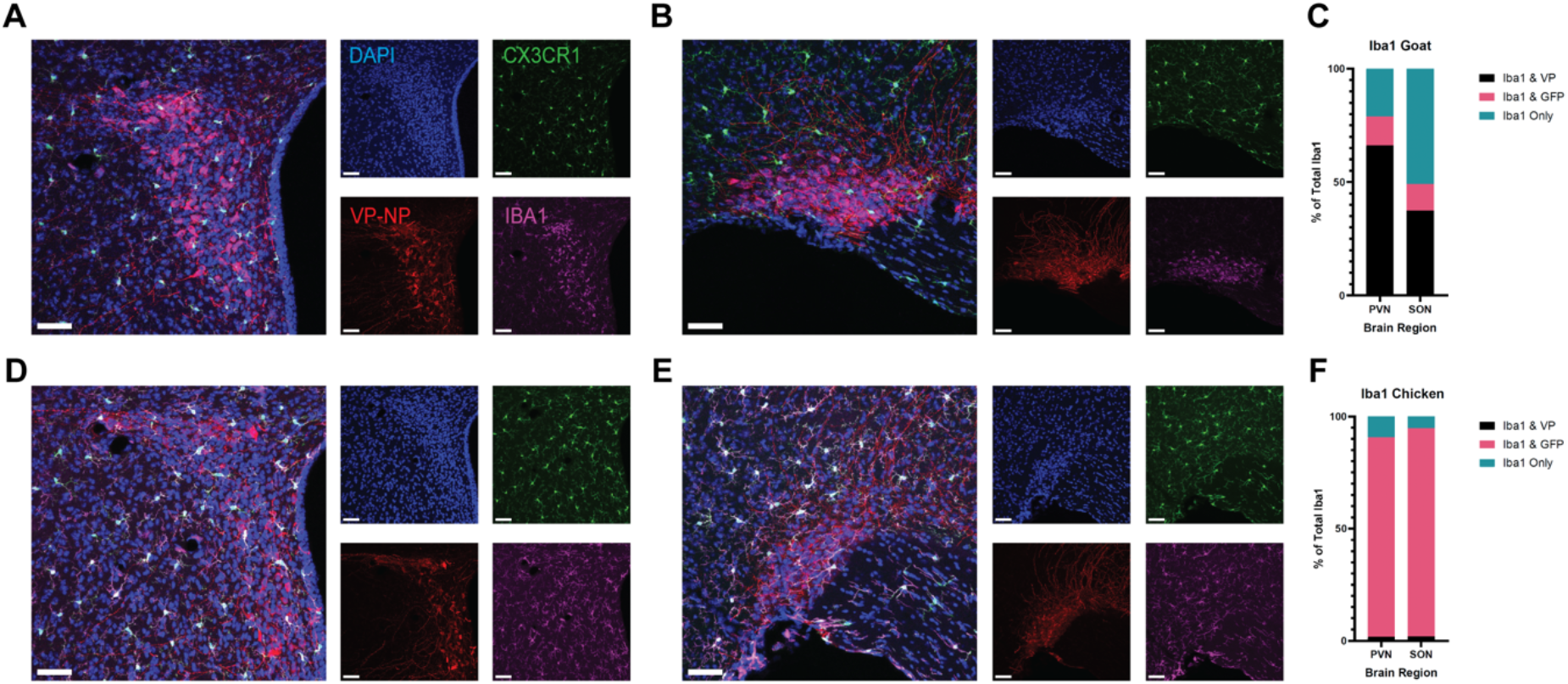
When compared to Iba1-goat, Iba1-chicken displays little to no colocalization with vasopressin, and high levels of colocalization with CX3CR1-GFP. A-B) Representative images of the PVN (A) and SON (B). C) Comparison of the percentage of cells labeled by Iba1-goat antibody alone, Iba1-goat colocalized with CX3CR1-GFP, and Iba1-goat colocalized with VP within the PVN and the SON. 66.2%±4.1 of Iba1+ cells in the PVN and 37.4%±3.1 of Iba1+ cells in the SON are co-labeled with Iba1-goat and VP. 12.7%±1.9 and 11.7%±2.5 of Iba1-goat cells co-labeled with CX3CR1-GFP in both the PVN and the SON, respectively, and the remaining 21.1%±4.5 of cells in the PVN and 50.9%±2.2 of the cells in the SON were labeled with only Iba1-goat. F) Comparison of the percentage of cells labeled by Iba1-chicken alone, Iba1-chicken colocalized with CX3CR1-GFP, and Iba1-chicken colocalized with VP within the PVN and the SON. 1.8%±0.9 and 2.0%±1.8 of the Iba1+ cells labeled by Iba1-chicken colocalized with VP in either the PVN or SON, respectively. 88.8%±6.1 of the Ibal+ cells in the PVN and 92.7%±5.2 of the cells in the SON were co-labeled with Iba1-chicken and CX3CR1-GFP. The remainder of Iba1+ cells in the PVN (9.7%±5.0) and the SON (5.3%±5.0) were labeled with only Iba1-chicken. Data are Mean ± SEM. Scale bar = 20 μm, blue = DAPI, green = CX3CR1-GFP, red = VP, magenta = Iba1-goat (A-B) or Iba1-chicken (D-E).

**Figure 5.**
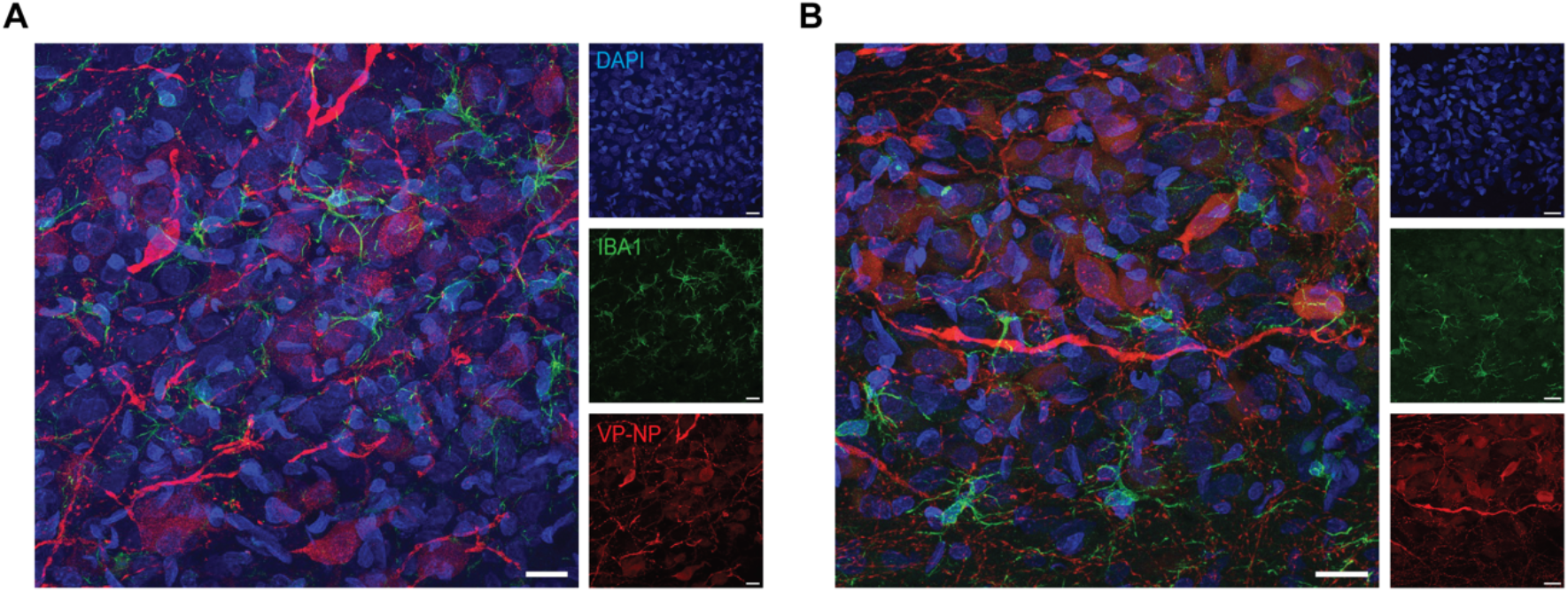
In Wistar rats, Iba1-goat displays normal labeling of microglial cells. A-B) Representative images of the PVN (A) and SON (B). Scale bar = 20 μm, blue = DAPI, green = Iba1-goat, red = VP.

Our results indicate that it is important to validate antibody efficacy when using new antibodies prior to beginning experimental data collection in order to ensure that cell labeling is both specific and accurate. This concept is particularly important when using new antibodies for the same molecular target (in this case, anti-Iba1) made in different host species. While we did not observe that specific lots of anti-Iba1-goat produced these erroneous effects, it is still possible that these errors may arise in other antibodies in specific lots.

Overall, not all commercially available antibodies for anti-Iba1 detect microglial cells with specificity as advertised in C57BL/6 mice. Thus, researchers should use caution when employing new antibodies in experiments, validating that new antibodies function as expected in all regions of interest by using alternative markers for the cell type of interest that have already been validated by the lab. Furthermore, the results of this study emphasize the importance of validating all antibodies prior to experimentation, as this may not be the only case in which there is a species- and region-specific cross-reactivity of an antibody with another cell type.

## Acknowledgements

The authors would like to thank Dr. Matthew Kirchner for his assistance during these experiments, as well as Logan Ouellette and Sylvie Call for their technical assistance. The authors would also like to acknowledge the Imaging Core Facility at Georgia State University for their excellent support and assistance in this work, and the Georgia State University Division of Animal Resources for outstanding animal care.

---

**Extended Data Figure 1-1.**
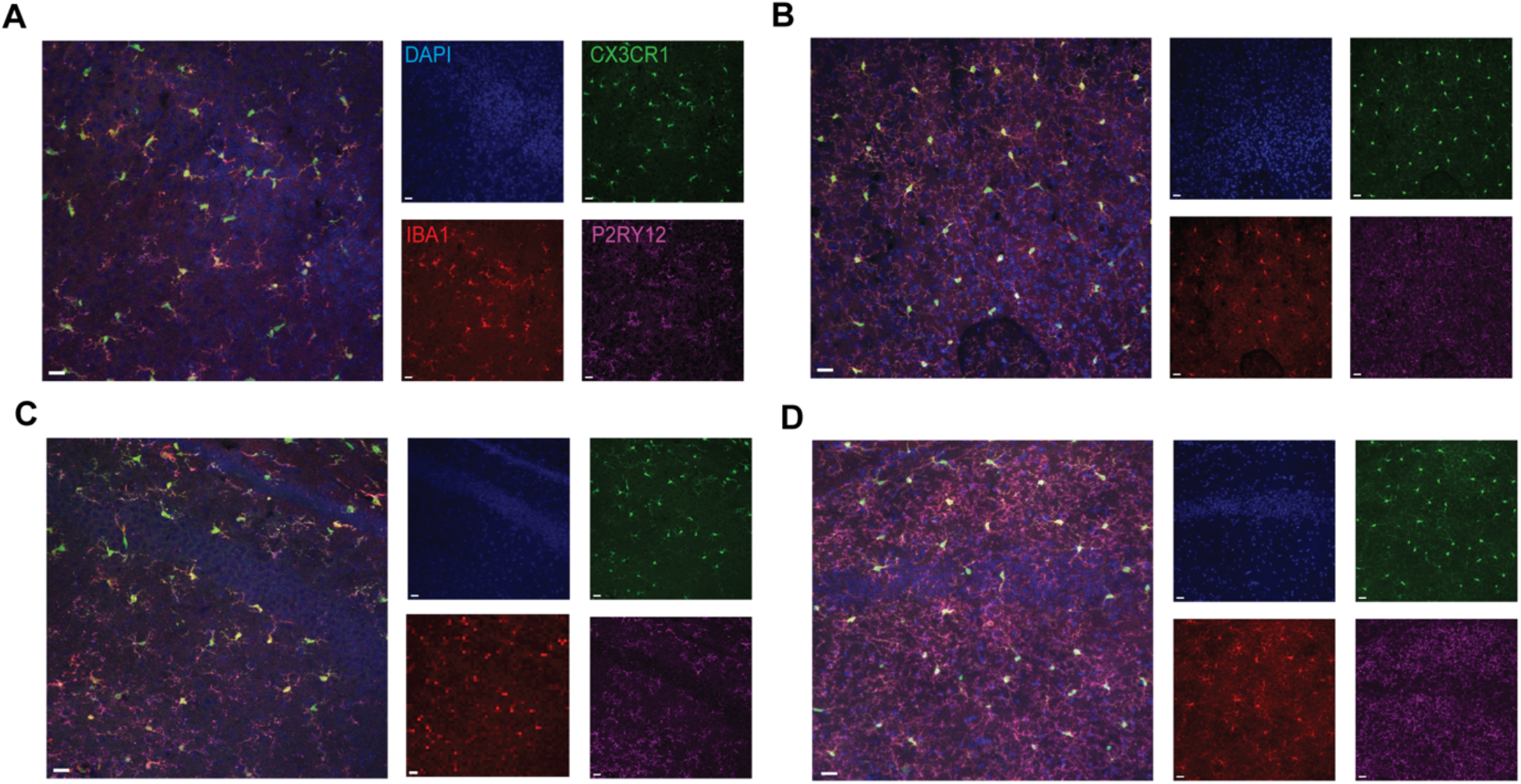
Excessive Iba1-goat staining is confined to specific regions of the mouse brain. A-B) Representative images of the parietal cortex in A) pups and B) adults. C-D) Representative images of the CA1 of the hippocampus in (C) pups and (D) adults. Scale bar = 20 μm, blue = DAPI, green = CX3CR1, red = Iba1-goat, magenta = P2RY12.

## Notes

<u>Conflict of Interest:</u> Authors report no conflict of interest.

<u>Funding Sources:</u> This work was supported by NIMH K99/R00 Pathway to Independence Award #MH120327, NARSAD Young Investigator Grant #31308 from the Brain & Behavior Research Foundation and The John and Polly Sparks Foundation, and Whitehall Foundation Grant #2022-08-051 awarded to JLB, as well as NIH #R01HL162575 to JES. Research reported in this publication was supported by the Office of The Director, National Institutes of Health and National Institute of General Medical Sciences under Award Number S10OD032336-01. The content is solely the responsibility of the authors and does not necessarily represent the official views of the National Institutes of Health.

### Competing Interest Statement

The authors have declared no competing interest.

